# Cell types associated with human brain functional connectomes and their implications in psychiatric diseases

**DOI:** 10.1101/2024.12.11.627878

**Authors:** Pengxing Nie, Yafeng Zhan, Renrui Chen, Ruicheng Qi, Cirong Liu, Guang-Zhong Wang

**Affiliations:** CAS Key Laboratory of Computational Biology, Shanghai Institute of Nutrition and Health, University of Chinese Academy of Sciences, Chinese Academy of Sciences, Shanghai 200031, China; Institute of Neuroscience, CAS Center for Excellence in Brain Science and Intelligence Technology, Chinese Academy of Sciences, Shanghai, 200031, China

**Author notes:** These authors contributed equally (Pengxing Nie, Yafeng Zhan).

**Keywords:** Brain cell type, functional connectome, resting-state functional network, neuropsychiatric disease, the cell-cell communication

## Abstract

Cell types are fundamental to the functional organization of the human brain, yet the specific cell clusters contributing to functional connectomes remain unclear. Using human whole-brain single-cell RNA sequencing data, we investigated the relationship between cortical cell cluster distribution and functional connectomes. Our analysis identified dozens of cell clusters significantly associated with resting-state network connectivity, with excitatory neurons predominantly driving positive correlations and inhibitory neurons driving negative correlations. Many of these cell clusters are also conserved in macaques. Notably, functional network connectivity is predicted by cellular communication among these clusters. We further identified cell clusters linked to various neuropsychiatric disorders, with several clusters implicated in multiple conditions. Comparative analysis of schizophrenia and autism spectrum disorder revealed distinct expression patterns, highlighting disease-specific cellular mechanisms. These findings underscore the critical role of specific cell clusters in shaping functional connectomes and their implications for neuropsychiatric diseases.

---

Cellular typing has provided profound insights into the cellular composition of the human brain^1–5^. Through single-cell RNA sequencing, variations in gene expression, functionality, and disease-related changes across different human cell types can be revealed^1, 6–10^. Similarly, comparable studies have been conducted in other primates, including rhesus monkeys^11^, macaques^12–14^, and marmosets^13, 15, 16^. These studies have uncovered genetic differences among neurons, astrocytes, oligodendrocytes, and other cell types across various brain regions. Additionally, an increasing amount of single-cell data^17, 18^, including those related to psychiatric diseases, is being integrated, providing the scientific community with valuable resources.

Functional imaging, which measures brain activity by detecting changes in blood oxygen level-dependent (BOLD) signals, has been employed in studies of numerous cognitive tasks and to investigate brain evolution across species^19–24^. The integration of human brain transcriptomic data, such as that from the Allen Human Brain Atlas^25^, with functional imaging has enhanced our understanding of the molecular basis of brain function^26–31^. This approach deepens our insights into essential biological process, neurological disease mechanisms, and therapeutic responses^32–36^. Additionally, combining functional imaging with transcriptomic analyses allows for the identification of brain regions and key gene networks involved in regulating cognitive activity^28,31, 32^.

Since brain cell types are the fundamental units of brain function, it is crucial to explore which cell types are involved in the functional connectome. Previous studies have primarily focused on correlating resting-state imaging signals with bulk transcriptomic data, while research on the role of cell clusters in functional connectomes at the single-cell level across the entire cortex remains limited. Here, we integrated high-resolution human brain single-cell atlases with resting-state and task-based MRI data to investigate how different cell clusters affect imaging signals and whether these clusters are linked to the pathology of neuropsychiatric diseases. We also obtained whole-brain spatial transcriptomic and functional imaging data from macaques to conduct an evolutionary conservation analysis of this relationship. Furthermore, we examined the role of functional signal-associated cell clusters and genes in neuropsychiatric diseases, including schizophrenia and autism spectrum disorder. Our study significantly enhances the crosstalk between single-cell transcriptomics and functional connectomes, revealing the importance of cellular-level heterogeneity in human neuropsychiatric disorders.

## Results

### Integrated analysis reveals contributions of cell clusters and genes to human resting-state functional networks

To investigate the contributions of different cell clusters to resting-state functional networks (rsf-networks), we conducted an integrated analysis combining single-cell RNA sequencing data^1^ from multiple regions of the human brain with resting-state functional magnetic resonance imaging (rsf-MRI) data from the Human Connectome Project (HCP). First, we constructed a rsf-network comprising 49 cortical regions (Fig. 1a). Within this network, we calculated the connectivity strength of each region with all other regions (Fig. 1a), revealing that ventral and dorsal brain regions exhibit higher connectivity^37, 38^, while primary visual areas display lower connectivity^39,40^. Next, using human whole-brain single-cell sequencing data, we calculated the proportions of various cell clusters across 28 cortical regions (Supplementary Table 1). We then applied Partial Least Squares (PLS) regression to explore the relationship between the proportions of different cell clusters and rsf-network connectivity strength. Statistical significance for each component was assessed using permutation tests (10,000 times), which showed that the first PLS component (PLS1) accounted for the maximum covariance between cell cluster proportions and connectivity (p = 0.0009) (Fig. 1b). Our analysis revealed a strong positive correlation between the spatial distribution of the PLS1 scores and functional connectivity strength (r = 0.82, *p* = 9.3 × 10⁻⁸) (Fig. 1c), suggesting that the composition of different cell types plays a crucial role in influencing rsf-network connectivity.

**Fig. 1:**
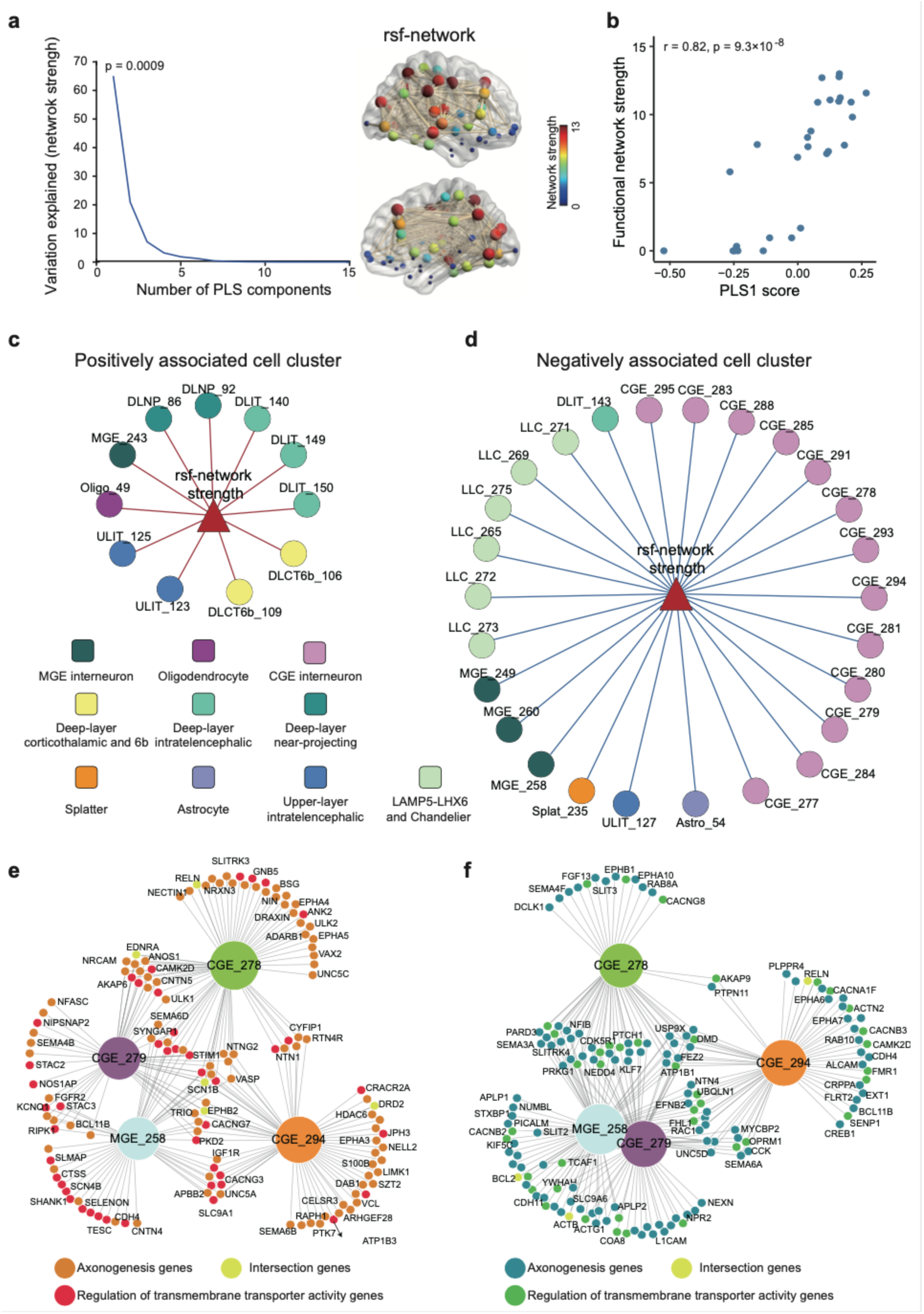
Identification of rsf-network-associated cell clusters and genes in the human cortex. **a**, Percentage of variance explained by different principal components in the PLS analysis of the relationship between rsf-network strength and the proportion of cell clusters across various brain regions. The inset panel displays the rsf-network of 49 brain regions in the human brain, with node size and color representing connectivity strength. **b**, Correlation between rsf-network strength and the PLS1 score. **c**. Cell clusters positively associated with rsf-network strength. **d**. Cell clusters negatively associated with rsf-network strength. Lines indicate significant correlations, with red representing positive and blue representing negative associations. **e, f,** Genes identified in cell clusters CGE_278, CGE_279, CGE_294, and MGE_258 that are positively (**e**) or negatively (**f**) correlated with rsf-network strength, and their roles in axonogenesis and the regulation of transmembrane transport protein activity.

Ultimately, we identified 11 cell clusters positively correlated with the connectivity of rsf-networks (Figure 1A) and 26 cell clusters negatively correlated with this connectivity (Figure 1B and Supplementary Table 2). Among the positively correlated cell clusters, 9 were excitatory neurons, and only one was an inhibitory neuron, a specific subtype of SST neurons (Fig. 1d). Conversely, of the 26 negatively correlated cell clusters, 23 were inhibitory neurons (Fig. 1e). To verify the robustness of our analysis, we varied the correlation threshold used to construct the rsf-networks. We found that when the correlation threshold ranged from 0.15 to 0.4, the composition of cell clusters across brain regions remained highly correlated with functional network connectivity strength (correlation coefficient > 0.8; Extended Data Fig. 1a). Furthermore, across different thresholds, the overlap of identified cell clusters associated with connectivity consistently remained above 70% (Extended Data Fig.1c). Additionally, by systematically removing one brain region at a time and performing the analysis on the remaining data, we found that over 75% of the associated cell clusters were consistently identified across all experiments (Extended Data Fig. 2). Our findings highlight the importance of different cell clusters in rsf-networks, emphasizing the distinct roles that excitatory and inhibitory neurons play in network connectivity.

To investigate the molecular mechanisms underlying rsf-networks, we generated a pseudobulk gene expression matrix for each cell cluster in each brain region. Using Partial Least Squares (PLS) analysis, we identified genes specifically associated with connectivity strength within cell clusters significantly correlated with these functional networks. Our results showed that many of these connectivity-associated genes were found exclusively within specific cell clusters (Extended Data Fig. 3a,b). The ion transport-related genes *CABP1* and *SCN1B* were identified as positively correlated with functional network strength in 22 cell clusters (Supplementary Table 3). In contrast, the calcium-binding protein-encoding gene *CRACR2A* was uniquely identified as positively correlated with functional network strength in the CGE_294 cell cluster. Furthermore, the gene *CAMK2D* exhibited a positive correlation with network strength in seven interneuron cell clusters (CGE_278, CGE_279, CGE_283, CGE_284, CGE_285, CGE_291, MGE_260) and a negative correlation in another seven cell clusters (Astro_54, CGE_293, CGE_294, CGE_295, LLC_271, MGE_243, MGE_249), suggesting that *CAMK2D* may play distinct roles across different neuronal subtypes. Next, we performed functional enrichment analysis on the genes positively and negatively associated with network connectivity for each cell cluster. We found that in interneuron cell clusters CGE_279 and MGE_258, the genes positively associated with connectivity were mainly involved in ion transport (Fig. 1f, Extended Data Fig. 4, adjusted p < 0.01). In contrast, in interneuron cell clusters CGE_278 and CGE_294, these genes were primarily involved in axon growth and development (Fig. 1f, Extended Data Fig. 4, adjusted p < 0.05). Conversely, the genes negatively associated with network connectivity were predominantly involved in neuronal projection, neurogenesis, neuronal development, and differentiation (Extended Data Fig. 5, adjusted p <0.05). Our findings highlight the diverse molecular mechanisms underlying rsf-networks, revealing distinct gene contributions to connectivity.

### Many cell clusters associated with rsf-networks are also involved in task-related brain states

To determine whether the cell clusters associated with rsf-networks also play roles in task-related brain states, we obtained functional imaging data for seven tasks from the Human Connectome Project (HCP). These tasks included working memory, motor function, gambling, language processing, social cognition, emotion processing and relational processing. A significant positive correlation was observed between the regional connectivity of resting-state networks and the connectivity during working memory, motor, and gambling tasks (r > 0.38, p < 0.05) (Extended Data Fig. 6a)^41, 42^. In contrast, a negative correlation was found between resting-state network connectivity and connectivity during language processing and social cognition tasks tasks (r < - 0.4, p < 0.004) (Extended Data Fig. 6a), consistent with previous reports^43, 44^. For the cell clusters significantly associated with resting-state network connectivity, many also showed strong correlations with the activity of task-related functional networks. Specifically, for the working memory task, four interneuron cell clusters, CGE_293, CGE_279, CGE_285, and MGE_258, were negatively correlated with task activation (r > 0.38, p < 0.05, Extended Data Fig. c, supplementary table 4). Among these, CGE_293, CGE_279, and CGE_285 are subtypes of VIP (vasoactive intestinal peptide) neurons, while MGE_258 is a subtype of PVALB (parvalbumin) neurons. In the motor task, two inhibitory neuron cell clusters, DLNP_86 (a deep-layer near-projection neuron) and CGE_295 (a VIP neuron), showed significant correlations (r > 0.4, p < 0.03, Extended Data Fig. b,c). For language processing and social cognition tasks, most of the correlated cell clusters overlapped (seven overlapped cell clusters, Extended Data Fig. 7b, c, Supplementary Table 4), indicating that these two cognitive functions may share common neural foundations. These findings provide new insights into how the brain processes information under different task conditions, highlighting the complex interactions between cell clusters, resting-state networks, and task-related activities^41, 45^.

### The relationship between rsf-networks and cell clusters is conserved between humans and macaques

To determine whether the relationship between rsf-networks and cell clusters is conserved across species, we utilized single-cell spatial transcriptomic data from macaques^12^, covering 143 coronal regions spanning the entire left cortical hemisphere. Additionally, we obtained two sets of macaque resting-state imaging data, each subdivided into 66 functional regions based on the D99 macague atlas. We constructed rsf-networks for macaques using both datasets and applied the same analytical workflow used for the human data to ensure comparability. Our findings revealed a significant correlation between the network connectivity of macaque rsf-networks and the cellular composition of distinct brain regions (r > 0.5, *p* < 0.05 in both datasets, Extended Data Fig. 7), which is consistent with our observations in the human brain. Furthermore, we identified a total of seven cell clusters conserved between humans and macaques that are associated with connectivity in rsf-networks. Among these, four cell clusters (MGE_243, ULIT_125, ULIT_123, and DLIT_149) were positively correlated with network connectivity, while three (CGE_279, CGE_288, and DLIT_143) were negatively correlated. Notably, many of these cell clusters were located in layers L2 and L3 (Extended Data Fig. 8), suggesting that these layers may play a key role in the formation and maintenance of functional networks in resting-state conditions.

### Cell-cell communication among rsf-network cell clusters predicts functional network connectivity

We hypothesized that rsf-networks are related to the communication between these rsf-network-associated cell clusters. To test this hypothesis, we constructed brain region communication networks based on the communication strength between cell clusters across different brain regions, including clusters that are positively correlated, negatively correlated, and uncorrelated with network connectivity. Interestingly, we found a significant positive correlation between the cellular communication network formed by positively correlated cell clusters and the connectivity of the rsf-network (r = 0.35, *p* = 3.5 × 10^!"#^; Fig. 2a). Notably, a similar correlation was found when analyzing rsf-network strength and the connectivity of the cell-cell communication network (r = 0.42, *p* =0.03, Fig. 2b). We further estimated the communication strength among cell clusters within each brain region and found that the negatively correlated cell clusters exhibited the highest cell-cell communication strength in most brain regions (Fig. 2c, *p <* 1 × 10^!$^,Extended Data Fig. 9). This indicates that these rsf-network negatively correlated cell clusters primarily influence rsf-network locally. Together, these results suggest that cellular communication of cell clusters correlated with the rsf-network may play a key role in the formation and maintenance of these functional networks.

**Fig. 2:**
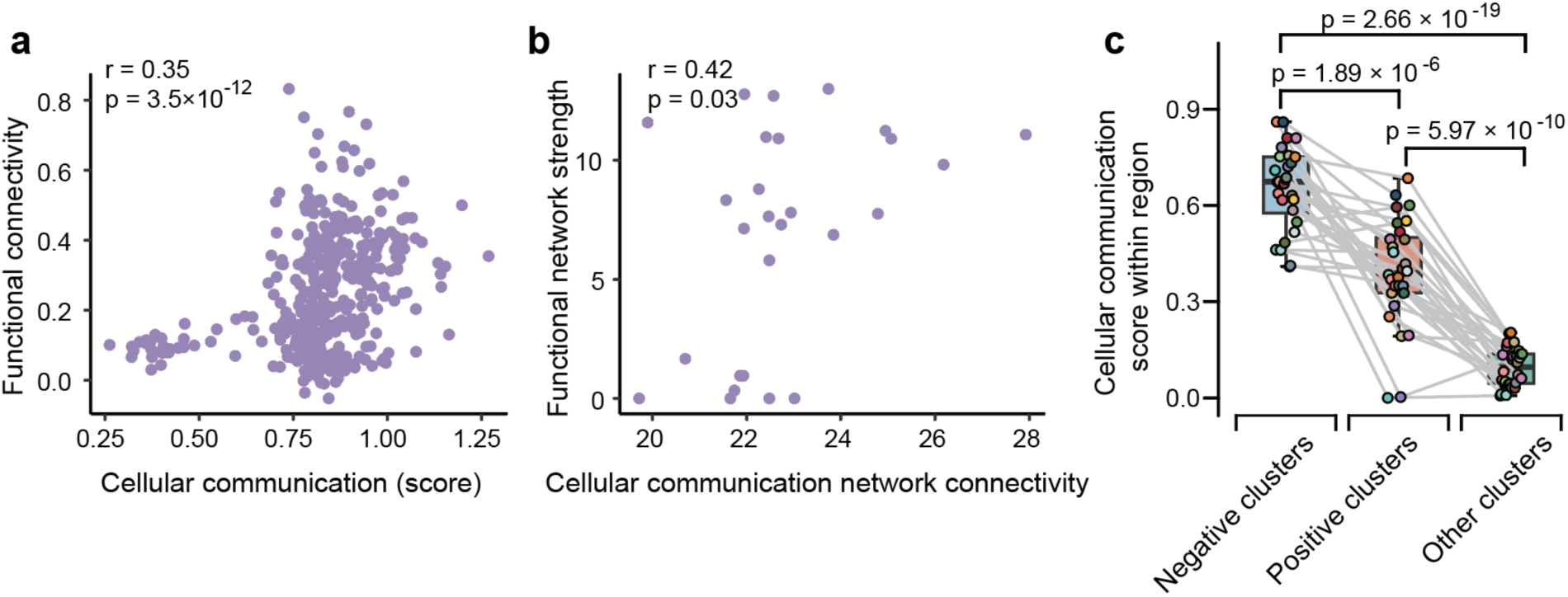
The rsf-network connectivity is strongly correlated with the human brain’s cellular communication network. **a**, Correlation between the interregional cellular communication score (weight) constructed from positively associated cell clusters, and the rsf-network connectivity. **b**. Correlation between interregional cellular communication connectivity constructed from positively associated cell clusters and the rsf-network strength. **c**. Comparison of cellular communication score (weight) among negatively associated, positively associated, and other cell clusters within each brain region. Colored points represent individual brain regions. Paired t-tests were used to assess statistical significance.

### Human rsf-network cell clusters involved in neuropsychiatric disorders and memory activity

To investigate whether genes significantly associated with rsf-networks in human brain cell clusters are linked to neuropsychiatric disorders, we obtained gene sets related to these diseases from related research^46^. The diseases considered include autism spectrum disorder (ASD), Bipolar Disorder (BD), Depression, and Schizophrenia (SCZ). We then conducted an in-depth analysis of the potential connections between these disease risk genes and the rsf-network-related genes we identified. Our findings revealed that, in multiple cell clusters, genes associated with rsf-network connectivity are significantly related to psychiatric diseases (Fig. 3a). For instance, schizophrenia-related genes showed significant enrichment among the connectivity-associated genes in 18 cell clusters (adjusted p < 0.02). These include two deep-layer intratelencephalic cell clusters (DLIT_149 and DLIT_150) and 16 inhibitory cell clusters, specifically, one LAMP5-LHX6 and Chandelier cell cluster, three MGE interneuron clusters, and 12 CGE interneuron clusters. Similarly, autism-related genes demonstrated significant enrichment in 13 cell clusters (adjusted p < 0.05), comprising one deep-layer intratelencephalic cell cluster (DLIT_149) and 12 inhibitory cell clusters (three MGE interneurons and nine CGE interneurons). Bipolar disorder-related genes showed significant enrichment in 19 cell clusters clusters (adjusted p < 0.034), including two deep-layer intratelencephalic cell clusters (DLIT_149 and DLIT_150) and 17 inhibitory cell clusters (one LAMP5-LHX6 and Chandelier cluster, four MGE interneurons, and 12 CGE interneurons).

**Fig. 3:**
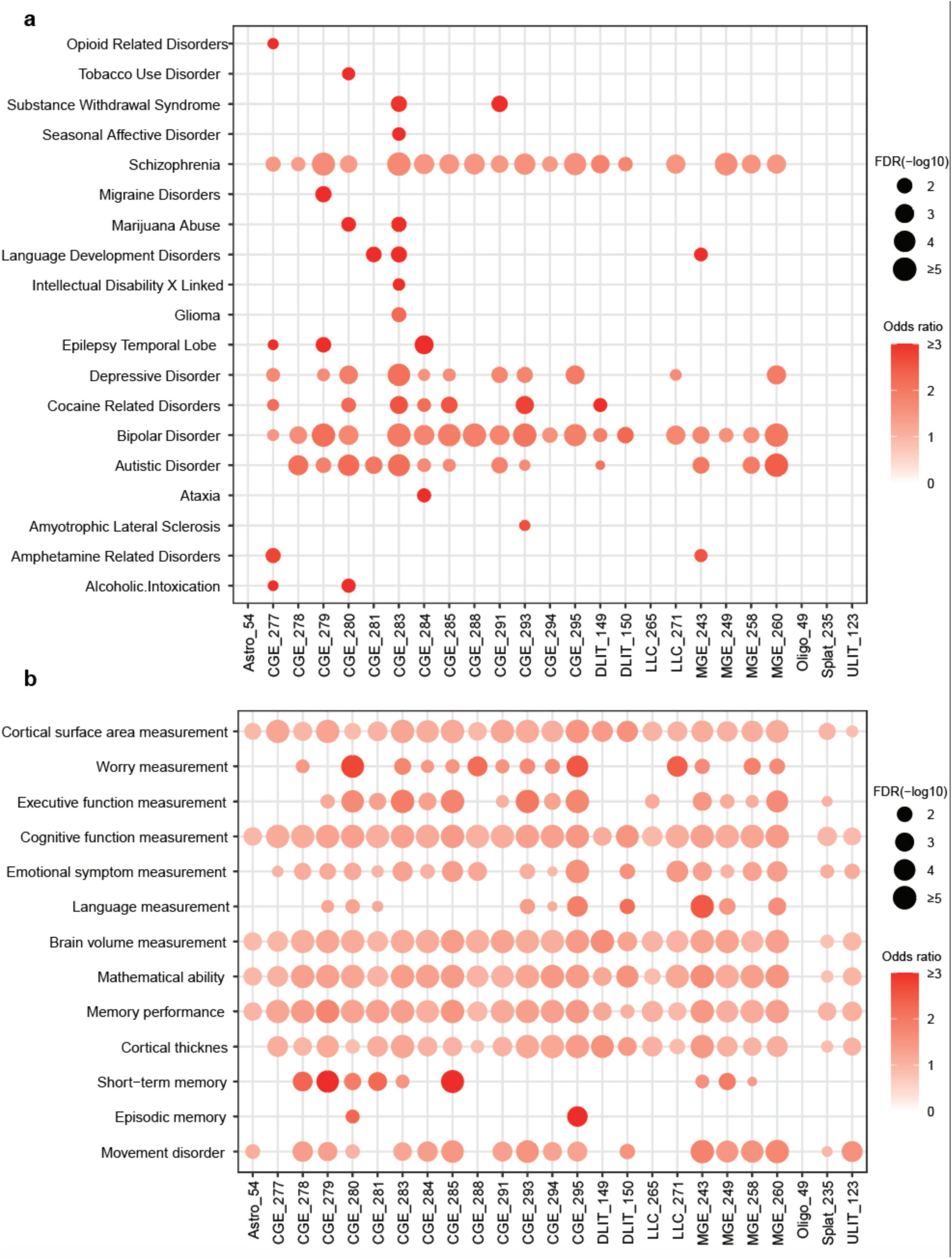
Relationship between rsf-network–associated genes and neuropsychiatric disorders as well as brain cognitive phenotypes. **a**, Enrichment of rsf-network–associated genes in each cell cluster for neuropsychiatric disorders. **b**, Enrichment of rsf-network–associated genes in each cell cluster for brain functions. The size of each bubble reflects the statistical significance of the enrichment, with larger bubbles indicating greater significance. The color intensity of each bubble represents the effect size of the association, measured as the Odds Ratio (OR).

Notably, the genes associated with rsf-network connectivity in interneuron cell clusters MGE_260, CGE_279, CGE_280, CGE_283, CGE_284, CGE_285, CGE_291, and CGE_293 are mostly linked to schizophrenia, autism, bipolar disorder, and depression. Furthermore, we found that the rsf-network connectivity-associated genes in CGE interneuron cell cluster CGE_283 are related to the largest number of neurological diseases, totaling 11 different disease types. Following this, CGE interneuron clusters CGE_277 and CGE_280 are associated with eight types of neuropsychiatric diseases. We also observed that rsf-network-associated genes are linked to certain neurological diseases only in some specific cell clusters, such as epilepsy, language development disorders, dementia, and alcoholism. These findings suggest that different neurological diseases may affect specific neuronal populations, revealing the cellular heterogeneity underlying these diseases.

Our analysis also shows that rsf-network-associated genes in almost all cell clusters are significantly linked to brain structures such as cortical surface area, cortical thickness, and brain volume (Fig. 3b). This suggests that these genes not only influence brain function at the molecular level but may also affect brain structure on a macroscopic scale. Additionally, rsf-network-associated genes are generally associated with genes related to various cognitive functions, emotions, mathematical abilities, and memory performance (Figure 3B). In many cell clusters, their association with short-term memory and episodic memory is also observed. This result has been validated using another set of memory encoding data^47^ (Extended Data Fig. 10). This may reflect the highly specific neural basis of these memory processes.

The cortical region and cerebellum collaborate in many advanced functions^48^. For instance, in certain neuropsychiatric disorders, the functional gradient between the brain and cerebellum may be altered^49–51^. Our recent study identified functional gradients reflecting the anterior-posterior connectivity of the cerebellum in resting-state fMRI data across multiple species, and identified the associated functional gradient-related genes^16^. Here, we found that many genes significantly associated with cortical resting-state fMRI signals were also linked to cerebellar functional gradients in macaques (odds ratio range = 1.22∼1.68, adjusted p < 0.03) and marmosets (odds ratio = 1.25∼1.60, adjusted p < 0.04, Extended Data Fig. 11), while the opposite is observed in mice. This finding suggests potential molecular similarities underlying functional connectivity across different brain regions in primates, while also highlighting species-specific differences in cerebellar connectivity and gene expression.

### Genes associations with rsf-networks are differentially expressed in schizophrenia and autism patients

We found that in both autism and schizophrenia, genes whose associations with rsf-networks are altered are significantly enriched among the rsf-network genes we identified (Extended Data Fig. 12). To further investigate the possible mechanisms linking these genes to neuropsychiatric disorders at the single-cell level, we obtained single-cell datasets of the prefrontal cortex from patients with schizophrenia and autism spectrum disorder, as well as from healthy controls^52, 53^. We conducted single-cell level analyses on the genes positively and negatively associated with rsf-networks across all rsf-network cell clusters. The results showed that these genes exhibited significant differential expression across various cell types between healthy individuals and patients. Specifically, in schizophrenia patients, genes positively linked to rsf-network connectivity were highly expressed in most neuronal cell types, while negatively associated genes were expressed at lower levels (Fig. 4a). In contrast, autism patients showed lower expression of positively associated genes in most neurons, and higher expression of negatively associated genes (Fig. 4b). This result may reflect different mechanisms of neural regulation in these two diseases, suggesting that the functions of these genes may vary under different pathological context.

**Fig. 4:**
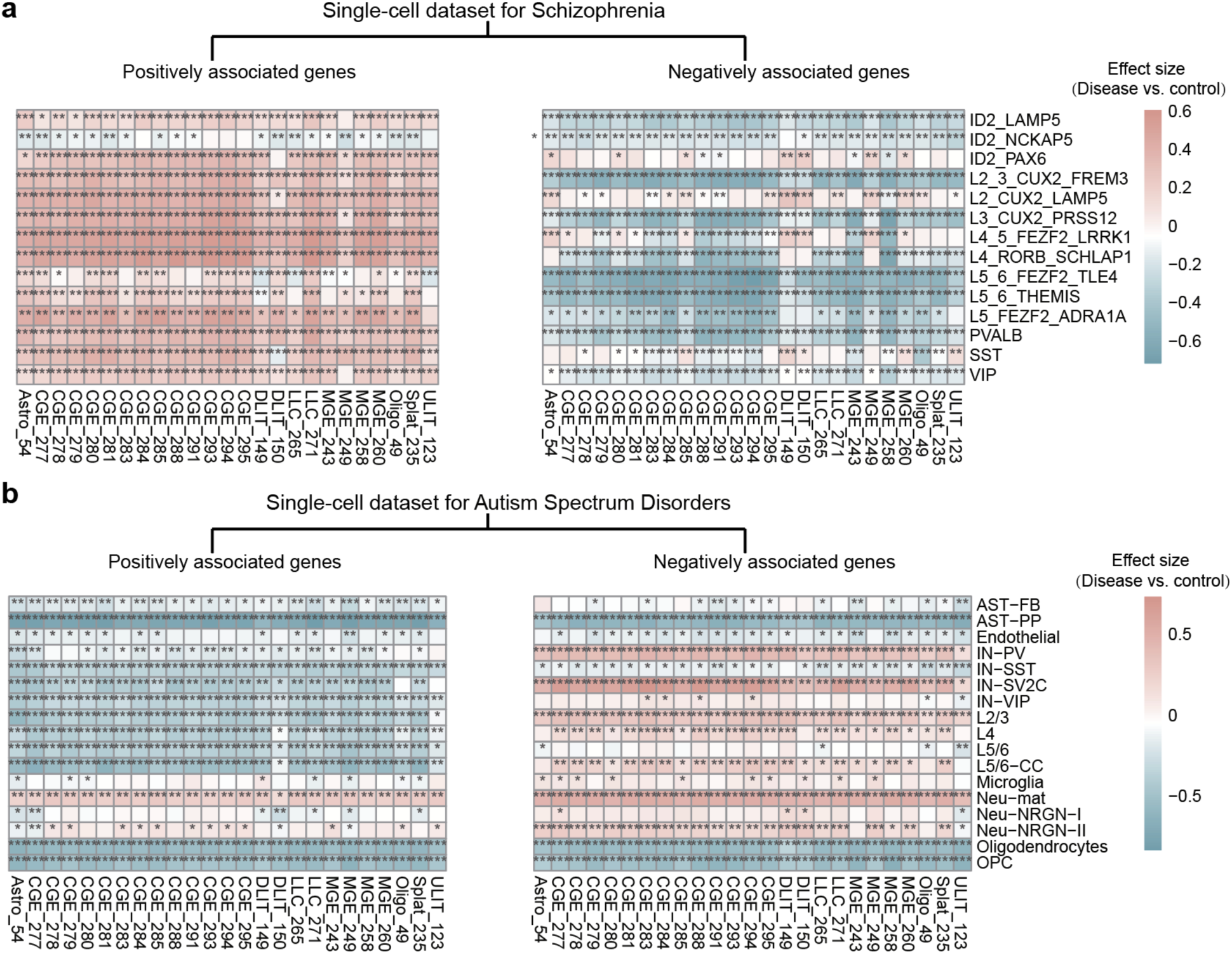
Differential expression of rsf-network cell clusters and genes in schizophrenia and autism spectrum disorder. **a**, Differences in the expression levels of rsf-network positively associated genes (left) and negatively associated genes (right) in each cell cluster, comparing individuals with schizophrenia and healthy controls. **b**, Differences in the expression levels of rsf-network positively associated genes (left) and negatively associated genes (right) in each cell cluster, comparing individuals with autism spectrum disorder and healthy controls. The color of each square (red or blue) represents the direction and magnitude of the effect size from t-tests, and asterisks denote statistical significance (* p < 0.05, ** p < 0.0001, *** p < 1 × 10^!"%^).

To validate these observed differences in rsf-network genes between schizophrenia and ASD, we analyzed single-cell sequencing data from 388 individuals, as reported by Prashant S. Emani et al^54^. This dataset included differentially expressed genes in patients versus healthy controls for both disorders. Enrichment analyses showed that rsf-network positively associated genes in various cell clusters were predominantly enriched among upregulated genes in schizophrenia (Extended Data Fig. 13). In contrast, in ASD, positively associated genes were mainly enriched among downregulated genes, whereas negatively associated genes were predominantly enriched among upregulated genes (Extended Data Fig. 13). These results further support the disease-specific molecular distinctions observed in rsf-network–associated genes.

Additionally, the observed low expression of both positively and negatively associated genes in non-neuronal cells underscores their potential involvement in autism spectrum disorders (Fig. 4b). These findings provide valuable insights into the molecular underpinnings of rsf-networks in autism and schizophrenia, emphasizing the cell-type-specific and disease-specific roles of these genes.

## Discussion

By integrating human whole-brain single-cell RNA sequencing data with resting-state functional magnetic resonance imaging (fMRI) data from the Human Connectome Project, we explored the relationship between cell cluster composition in 28 cortical regions and the connectivity of their functional networks. We identified cell clusters and associated genes linked to functional connectivity. Our results revealed that cell cluster composition significantly influences the connectivity characteristics of functional networks, with 11 cell clusters positively correlated and 26 cell clusters negatively correlated with rsf-network connectivity. Positively correlated clusters were primarily composed of excitatory neurons, while negatively correlated clusters were predominantly inhibitory neurons. Notably, the influence of cell clusters on functional network connectivity was largely mediated through cell-cell communication. Using a similar approach with functional imaging and spatial transcriptomic data from macaques, we found that cell cluster composition in different brain regions also significantly affects rsf-network connectivity. Moreover, many cell clusters associated with resting-state networks were also active during task performance, reinforcing the strong relationship between resting-state and task-based functional networks. Lastly, our study identified numerous cell clusters associated with various psychiatric disorders, underscoring their critical role in neuropsychiatric conditions^55^. By leveraging single-cell disease data, we uncovered significant differences in gene expression patterns between cell clusters in patients with schizophrenia and those with autism spectrum disorder (ASD), shedding light on distinct pathogenic mechanisms underlying these disorders.

Cell clusters with higher communication strength are more closely related to the connectivity of rsf-networks. This suggests that these associated cell clusters facilitate the formation and maintenance of rsf-networks by enhancing inter-regional crosstalk. This finding reveals the cellular basis of human brain functional networks from the perspective of cell communication. Consistent with our results, previous studies have shown that both excitatory and inhibitory neurons contribute to resting-state BOLD signals, but they play distinct roles^56–58^. While excitatory neurons play a dominant role in coordinating activity between different brain regions and network integration^59, 60^, inhibitory neurons contribute to the dynamics of resting-state networks by maintaining system stability and allowing the network to flexibly switch between different network states^61, 62^. Our results further demonstrate the fundamental role of cellular and molecular communication among excitatory neurons in higher cognitive functions^63^. Future studies could investigate the dynamic roles of communication between different cell types within functional networks, particularly under varying cognitive tasks, to gain a deeper understanding of the relationship between cellular activity and functional networks.

By examining the layer-specific distribution of rsf-network cell clusters across different brain regions in macaques, we found that conserved cell clusters are predominantly located in layers 2 (L2) and 3 (L3). Pyramidal excitatory neurons in these layers play a critical role in corticocortical interactions. These neurons are involved not only in local microcircuits but also in forming long-range connections between cortical areas, thereby facilitating the integration of information across distinct cortical regions^64–66^. Moreover, L2/3 neurons exhibit the ability to integrate excitatory and inhibitory inputs within cortical circuits^66, 67^. This input integration mechanism is crucial for maintaining the balance of cortical function, ensuring that appropriate neural activity is generated and finely tuned at specific times.

We also identified the potential roles of cell clusters associated with the connectivity of rsf-networks and found that various neuropsychiatric disorders (such as autism spectrum disorder, bipolar disorder, and schizophrenia) are significantly linked to genes within specific cell clusters^53, 68, 69^. This suggests that these disorders may either originate from or be influenced by particular neuronal subpopulations^68^. Notably, inhibitory neuron clusters, such as those from the MGE and CGE, are significantly enriched across multiple neuropsychiatric diseases, underscoring their crucial role in maintaining normal brain function. Genes within several cell clusters are associated with multiple neuropsychiatric disorders, indicating that these diseases may share common cellular mechanisms. For example, genes within the CGE_283 cell cluster are linked to more than 10 different neurological disorders, highlighting the central role of this cluster in various diseases. Targeting these key cell clusters could provide new therapeutic opportunities for these disorders.

Through the analysis of single-cell data from autism and schizophrenia, we discovered that the rsf-network gene sets showed significant differences between disease states and normal individuals. Interestingly, genes positively correlated with network connectivity were generally highly expressed in specific neuron clusters from schizophrenia patients, whereas these genes were underexpressed in ASD. This finding suggests that those neuron clusters in schizophrenia may be hyperactivated, leading to imbalanced neural network connectivity and dysfunctional integration across large-scale brain networks^70, 71^. In contrast, the gene expression patterns in ASD are associated with disrupted neural network^72, 73^. These significant differences highlight distinct neurocellular mechanisms underlying the two disorders.

Our study has several limitations. First, the brain cell cluster information is based on RNA expression data rather than single-cell proteomic data, which may more accurately reflect functional connectome signals. This limitation could be addressed in future studies through the identification of brain-wide single-cell proteomes. Additionally, we lacked real-time data on cell-to-cell communication, which is essential for validating our findings related to cellular interactions. Acquiring such data in the future would strengthen the validation of our results. Finally, the absence of human brain-wide spatial transcriptomic data limits our understanding of the roles different cortical layers play in the functional connectome and neuropsychiatric disorders. The generation of spatial transcriptomic datasets will provide a clearer insight into these aspects.

## Materials and Methods

### Processing of human brain-wide single-cell data

The human whole-brain single-cell data used in this study were obtained from Siletti et al., comprising single-cell profiles derived from 105 brain sections. Processed cortical data^1^ were downloaded from the CellxGene database. Subsequently, the data were further processed using Seurat v4.3.0.1^74^. First, we removed non-coding genes and retained cells with 500-9,000 detected genes, excluding cells with >5% mitochondrial content. For data from the Perirhinal gyrus (PRG) and Rostral gyrus (RoG) regions, we additionally removed cells whose metadata did not correspond to their respective regions. To calculate the proportion of cell clusters across different cortical areas, only clusters present in more than half of the cortical regions were retained. Additionally, the 20% of clusters with the lowest mean and variation in their proportions across cortical areas were excluded. Ultimately, 151 cell clusters encompassing the expression profiles of 1,033,759 cells were retained for downstream analysis.

### Human resting-state and task fMRI data acquisition and processing

We obtained resting-state fMRI (rsfMRI) data from 1,200 participants in the Human Connectome Project (HCP) database (https://db.humanconnectome.org). The data were preprocessed using the ICA-fixed procedure (FMRIB’s ICA-based X-noisifier). Detailed information about data acquisition, preprocessing, and denoising methods for HCP rsfMRI data has been reported previously^75, 76^. Briefly, the data were acquired using a customized 3T Siemens Skyra scanner, with a voxel size of 2 mm, providing high-resolution rsfMRI for all participants. The preprocessing pipeline for rsfMRI data included temporal smoothing, spatial distortion correction, head motion correction, registration to T1-weighted anatomical images, transformation to MNI space, and high-pass temporal filtering. The Allen Brain Atlas^25^ was used as the region of interest (ROI) for constructing functional networks.

We also analyzed task-based fMRI data from the HCP, which included seven task categories: working memory, motor function, gambling, language processing, social cognition, emotion processing, and relational processing. The specific task contrasts analyzed were as follows: 1). working memory: 2-back vs. 0-back contrast. 2). motor function: the average activation across five motor tasks (tapping the left or right index finger, squeezing the left or right toes, and moving the tongue). 3). gambling: punishment (monetary loss) vs. reward (monetary gain). 4). language processing: story comprehension (listening to narratives) vs. math problem-solving (arithmetic).

5). social cognition: TOM (watching geometric objects engaged in social interactions) vs. random movement (watching randomly moving geometric objects). 6). relational processing: relational (comparing dimensions to distinguish two pairs of objects) vs. matching (matching objects based on verb categories). 7). emotion processing: faces (determining which of two angry/fearful faces at the bottom of the screen matches a face at the top) vs. shapes (performing the same task using shapes instead of faces). Individual task activation maps were generated using cross-run FEAT analysis in FSL^77^ (https://fsl.fmrib.ox.ac.uk), and subject-level (Level 2) analyses were conducted to compute effect sizes (Cohen’s d). The task activation maps provided by the HCP were smoothed using a 4 mm Gaussian kernel and mapped onto the fsLR-32k CIFTI surface space. Each hemisphere contained 32,492 vertices, and multimodal surface matching was applied to ensure alignment^78^.

### Macaques resting-state fMRI data acquisition and processing

Resting-state macaque fMRI data were obtained from the Nonhuman Primate Data Exchange (PRIME-DE) consortium (251) (http://fcon_1000.projects.nitrc.org/indi/indiPRIME.html). The study included two datasets from awake macaques, which are from the National Institute of Mental Health and the Lyon Neuroscience Research Center. Both datasets were processed using an identical preprocessing pipelines with AFNI^79^, FSL^80^, and ANTs^16, 81^. Briefly, spike artifacts were removed with AFNI’s 3dDespike, and slice-timing and head-motion corrections were performed using AFNI’s 3dTshift and 3dvolreg, respectively. Afterward, linear and quadratic trends, motion parameters (and their derivatives), high-motion volumes, white matter and CSF signals, and frequencies outside 0.01–0.1 Hz were regressed out using AFNI’s 3dDeconvolve and 3dTproject. All preprocessed fMRI data were spatially registered to the macaque NMT-V2 template^82^ using AFNI’s antsRegistration.

### Processing of macaque single-cell spatial transcriptomic data

We utilized single-cell spatial transcriptomic data from the rhesus monkey cortex obtained via Stereo-seq as reported by Chen et al.^12^ In total, we analyzed 119 slice datasets originating from the same individual monkey. For each cell, we extracted its spatial coordinates and calculated its distribution within the cortical layers.

### Identification of homologous cell clusters between humans and macaques

We used single-cell sequencing data from both humans and macaques and employed MetaNeighbor v4.4^83^ to map human cell clusters to corresponding macaque data. The topHits function was used to identify matching cell types, with a threshold > 0.9.

### Construction of rsf-networks and networks strength calculation

For the human resting-state functional imaging data, we utilized Workbench Command tools to construct and evaluate functional connectivity networks^84^. First, we extracted signal intensity from different brain regions using the wb_command -cifti-parcellate command on preprocessed imaging data. Next, we averaged the imaging signal across all runs and participants using the wb_command -cifti-average command. Finally, we computed correlations between brain regions with the wb_command -cifti-correlation command to construct the human brain functional network. For macaque resting-state functional imaging data, we used AFNI’s 3dNetCorr to compute functional connectivity for preprocessed macaque imaging data across brain regions^81^. To calculate functional network strength for human and macaque rsf-networks, we retained only connections with a correlation greater than 0.3. We then computed the weighted connectivity strength for each brain region by summing the strengths of its connections with other regions. This measure was used to assess the functional importance of each brain region within the overall network.

### Identification of cell clusters and genes associated with functional network strength via PLS analysis

We performed partial least squares (PLS) analysis^85, 86^ using the plsregress function in MATLAB R2021a to examine the relationship between functional network strength and both cell proportions across brain regions and gene expression at the cell cluster level. Statistical significance for variance explained by different components was determined through permutation testing with 10,000 times. We then used a bootstrap approach (10,000 times) to estimate the standard errors of the PLS1 loadings for each cell cluster and gene. From these bootstrap results, we calculated Z-scores to identify cell clusters and genes that contribute significantly to PLS1 (the first principal component). Cell clusters or genes with Z-scores > 3 were identified as positively associated, while those with Z-scores < -3 were identified as negatively associated.

### Construction of regional cell-cell communication networks

First, we analyzed the communication strength among different cell clusters within each individual brain region using the ‘createNeuronChat‘ and ‘run_NeuronChat‘ functions (with parameter M = 100) from the NeuronChat v1.0.0 R package^87^. Next, for brain-regional cell communication networks, we applied the same procedure to all pairwise combinations of brain regions, thereby estimating the communication strength between cell clusters across different regions. Subsequently, we constructed regional communication networks by calculating the average communication strength for cell clusters that were positively correlated with functional network strength, negatively correlated with functional network strength, and region-specific. Finally, we used these regional communication networks to compute the node degree.

### Functional enrichment analysis of rsf-network genes

We performed gene set enrichment analysis using the R package clusterProfiler v4.8.2^88^ to explore the functions of genes associated with network strength in different cell clusters. KEGG pathway enrichment and biological process (BP) enrichment analyses were carried out using the enrichKEGG and enrichGO functions, respectively. P-values were adjusted using the Benjamini-Hochberg method, with a threshold set at < 0.05 for significance.

### Analysis of rsf-network genes in relation to neuropsychiatric disorders and brain function

We obtained gene lists associated with brain disorders, as well as those related to cerebellar functional gradients in macaques, marmosets, and mice, from previous studies^46, 89, 90^. Using Fisher’s exact test, we examined the relationships between functional network genes and neuropsychiatric disorders, as well as cerebellar functional gradients. Additionally, we retrieved genes related to brain structure and function from the GWAS Catalog (https://www.ebi.ac.uk/gwas/)^91^. These gene sets include: movement disorder (EFO_0004280), emotional symptom measurement (EFO_0007803), worry measurement (EFO_0009589), mathematical ability (EFO_0004875), episodic memory (EFO_0004333), short-term memory (EFO_0004335), memory performance (EFO_0004874), cognitive function measurement (EFO_0008354), language measurement (EFO_0007797), executive function measurement (EFO_0009332), brain volume measurement (EFO_0006930), cortical surface area measurement (EFO_0010736), and cortical thickness (EFO_0004840).

### Differential analysis of rsf-network genes in schizophrenia and autism spectrum disorder single-cell data

We obtained preprocessed single-cell datasets for schizophrenia and autism spectrum disorder (ASD), along with corresponding control data, from studies by Mykhailo et al.^52^ and Velmeshev et al.^53^. Using the AddModuleScore_Ucell function from the R package Ucell v2.4.0^92^, we computed scores for rsf-network genes across various cell clusters in the single-cell data for these psychiatric disorders. All other parameters were set to their default values. T-tests were conducted to evaluate the significance of differences in expression levels between disease and control groups within specific cell clusters.

### Statistical analysis

Unless otherwise specified, all statistical analyses in this study were conducted using R v4.3.1. Gene set enrichment analysis was performed using two-sided Fisher’s exact tests. Multiple testing corrections were applied using the false discovery rate (FDR) method.

### Author contributions

G-Z. W. and C.L. supervised the study; P.N. and Y.Z. performed the data analysis with the help of R.Q. and R.C.; P.N. drafted the figures and the manuscript; P.N., G-Z.W. and C.L. wrote and edited the manuscript.

